# The TMEM16A Channel Mediates the fast polyspermy block in *Xenopus Laevis*

**DOI:** 10.1101/270629

**Authors:** Katherine L. Wozniak, Wesley A. Phelps, Maiwase Tembo, Miler T. Lee, Anne E. Carlson

**Author notes:** Correspondence (A.E.C.).

## Abstract

In externally fertilizing animals, such as sea urchins and frogs, prolonged depolarization of the egg immediately after fertilization inhibits the entry of additional sperm – a phenomenon known as the fast block to polyspermy. In the African clawed frog, *Xenopus laevis*, this depolarization is driven by a Ca^2+^-activated Cl^−^ efflux. Although the prominent Ca^2+^-activated Cl^−^ currents generated by immature *X. laevis* oocytes are conducted by xTMEM16A channels, little is known about which channels contribute to fertilization-competency in mature eggs. Moreover, the gamete undergoes a gross transformation as it matures from an immature oocyte into a fertilization-competent egg. Here we report the results of our approach to identify the Ca^2+^-activated Cl^−^ channel that triggers the fast block. Querying published proteomics and RNA-seq data, we identified two Ca^2+^-activated Cl^−^ channels expressed in fertilization-competent *X. laevis* eggs: xTMEM16A and xBEST2A. Furthermore, transcripts for these channels increase in abundance during gamete maturation. To determine if either of these mediates the fast block, we characterized exogenously expressed xTMEM16A and xBEST2A using pharmacologic inhibitors. None of the inhibitors tested blocked xBEST2A currents specifically. However, Ani9 and MONNA each reduced xTMEM16A currents by more than 70%, while only nominally inhibiting those generated by xBEST2A. Using whole-cell recordings during fertilization, we found that Ani9 and MONNA effectively diminished fertilization-evoked depolarizations. These results indicate that fertilization activates TMEM16A channels in *X. laevis* eggs and induces the earliest known event triggered by fertilization: the fast block to polyspermy.

**HIGHLIGHTS:**

- Protein for the channels xBEST2A and xTMEM16A is present in *X. laevis* eggs.

- The inhibitors MONNA and Ani9 effectively block xTMEM16A compared to xBEST2A.

- *Xenopus laevis* fertilization opens TMEM16A to trigger egg depolarization.

- The TMEM16A-mediated depolarization is critical for the fast block to polyspermy.

## INTRODUCTION

Fertilization of an egg by more than one sperm, a condition known as polyspermy, presents one of the earliest and most prevalent barriers to successful reproduction. In most sexually reproducing species polyspermy causes chromosomal abnormalities that are embryonic lethal [1]. Eggs have evolved multiple strategies to combat the entry of sperm into an already fertilized egg and to thereby avoid such catastrophic consequences [2]; however, the underlying molecular mechanisms are still poorly understood.

The two most common strategies for preventing polyspermy are the *fast block* and the *slow block* [3]. As their names imply, these mechanisms differ with respect to how quickly they occur. The fast block involves depolarization of the egg and occurs within seconds of fertilization [4]. Cross-species fertilization experiments demonstrated that sperm possess a voltage sensor that prevents their entry into a depolarized egg [5]; by a yet unknown mechanism, this voltage sensor allows sperm to detect whether an egg is depolarized and thus already fertilized. By contrast, the slow block involves the creation of a physical barrier surrounding the nascent zygote and takes several minutes to complete [1, 6]. Whereas the slow block occurs in all sexually reproducing species, the fast block is limited to externally fertilizing organisms, in which the sperm-to-egg ratio can be extremely high [4, 7, 8].

The fast block has been documented in diverse externally fertilizing organisms (*reviewed by* [9]), including fucoid algae [10], sea urchins [4], starfish [11], marine worms [12], and amphibians [7, 13]. For example, the African clawed frog *Xenopus laevis* is an externally fertilizing species that uses the fast block. The second messengers that trigger these fast blocks and the channels that conduct the depolarizing currents have not been identified in any species. Due to evolutionary distance and differences in habitat among these species, the precise mechanisms are likely to vary. Nevertheless, eggs capable of undergoing the fast block share three characteristics: their fertilization-preventing membrane depolarization, known as the *fertilization potential*, persists for one minute or more [4, 14] and is distinct from action potentials in other excitable cells such as neurons or cardiac myocytes [15]; when held at a depolarizing voltage of this kind, the eggs can be bound, but not entered, by sperm, even if unfertilized [4]; and when held at hyperpolarized potentials, the eggs can be fertilized by multiple sperm [4].

As in all frogs, the *X. laevis* fast block requires an increase of cytosolic Ca^2+^ and a depolarizing efflux of Cl^−^ [7, 14, 16, 17]; an event hypothesized to be mediated by a Ca^2+^-activated Cl^−^ channel (CaCC) [18, 19]. In eggs loaded with the Ca^2+^-chelator BAPTA, fertilization failed to evoke a depolarization or cleave the egg, thereby linking the absence of an electrical event with an absence of a developmental event [20]. Moreover, treating eggs with a Ca^2+^ ionophore, a lipid soluble compound that transports Ca^2+^ across the plasma membrane and increases intracellular [Ca^2+^], evoked a depolarization in the absence of fertilization [14]. The ionophore signaled depolarization demonstrated that increased intracellular Ca^2+^ is sufficient to trigger the fast block. A requirement for a Cl^−^ efflux was demonstrated by larger fertilization-evoked depolarizations recorded from eggs inseminated in low extracellular Cl^−^ and smaller depolarizations recorded from eggs inseminated in high extracellular Cl^−^ [14, 17]. Furthermore, replacing the dominant extracellular halide from Cl^−^ to Br^−^ or I^−^ led to no changes in membrane polarization or hyperpolarizations with fertilization, respectively [14]. Under these conditions, the magnitude and direction of the fertilization-evoked depolarization was directly linked to polyspermy. For example, multiple sperm penetrated all eggs inseminated in I^−^ compared to mostly monospermic inseminations in Cl^−^ [14]. Finally, insemination in Br_−_ resulted in an intermediate effect, with a mixture of monospermic and polyspermic embryos [14]. Together these experiments revealed both a prominent role for cytosolic Ca^2+^ increase and a Cl^−^ current in the fast block, and underscored the importance of a fertilization-evoked depolarization for ensuring monospermic fertilization. Here we sought to identify the Ca^2+^-activated Cl^−^ channel (CaCC) that mediates the fast block in *X. laevis*.

The channels expressed in the fertilization-competent *X. laevis* egg are not well studied, which is in stark contrast to the well-characterized channels found in the immature oocyte [e.g. 21]. Indeed, the oocytes and eggs of *X. laevis* are vastly different cells (Figure 1) [22]. Immature *oocytes* are located in the ovary, are arrested in prophase I, and cannot be fertilized. By contrast, *eggs* are located outside the *X. laevis* female (following ovulation and laying), are arrested in metaphse II, and are fertilization-competent (i.e. gametes). As the oocyte matures into an egg, many ion channels and transporters are internalized, including: Orai1, the pore-forming subunit of the store-operated Ca(2+) entry channel [23]; the plasma membrane Ca^2+^-ATPase (PMCA) [24]; and Na^+^/K^+^ ATPase [25]. In addition, oocyte maturation induces intracellular proteins that closely interact with the plasma membrane, including components of the cytoskeleton, to undergo transformations in their structural contacts [26]. Therefore, experimental findings regarding prominent CaCCs, namely TMEM16A [21], in *X. laevis* oocytes cannot be directly applied to eggs in the absence of further testing, and thus it was necessary to study the CaCCs in eggs directly.

**Figure 1.**
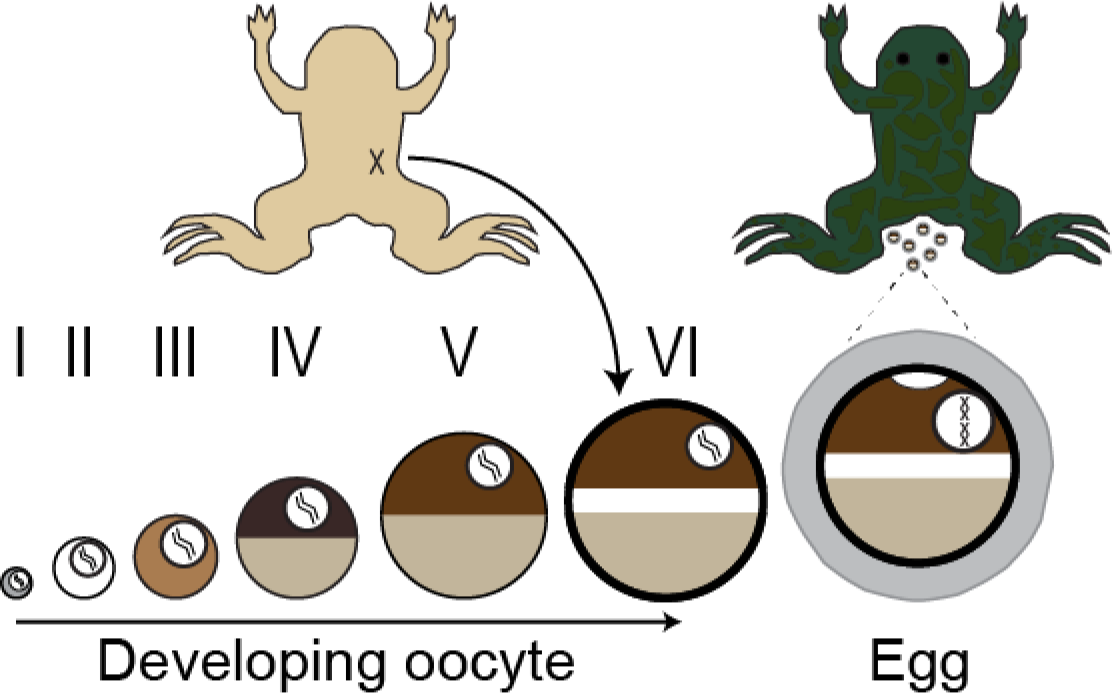
Schematic depiction of gamete development in female *X. laevis*. Immature oocytes, ranging from the youngest (stage I) to the most developed (stage VI), are located within the ovaries. These oocytes can be surgically removed from the abdomen of the frog (shown in ventral view at top left) and are commonly used by electrophysiologists. Upon hormonal induction, stage VI oocytes mature into fertilization-competent eggs, which are laid by the frog (shown in dorsal view at top right). Oocytes and eggs differ with respect to membrane-localized proteins as well as the structure of the cytoskeleton.

We sought to identify the channel that mediates the fast block in *X. laevis* eggs. Using existing proteomic and transcriptomic data from *X. laevis* oocytes and eggs [27, 28], we identified two candidate CaCCs: transmembrane protein 16 type a (TMEM16A) [21, 29, 30] and bestrophin 2a (BEST2A) [31, 32]. To distinguish between the currents produced by the *X. laevis* orthologs of these channels (xTMEM16A and xBEST2A), we exogenously expressed and pharmacologically characterized each. By applying this approach to whole-cell recordings of *X. laevis* eggs during fertilization, we demonstrate that it is xTMEM16A, and not xBEST2A, that produces the depolarizing current. Thus, we describe the first known ion channel that mediates the fast block.

## RESULTS

### Two candidate CaCCs accumulate in the egg and are candidates for the trigger of the fast block

To identify candidate CaCCs that may trigger the fast block in *X. laevis*, we interrogated two previously published high-throughput gene expression datasets. First, we examined the proteome of fertilization-competent eggs [24] and queried for all known ion channels (Figure S1 and Dataset S1). Three protein families containing CaCCs have been characterized to date: the CLCAs, the bestrophins (BEST), and the transmembrane protein 16s (TMEM16/ANO) [33]. We discovered that only one member of the BEST family, xBEST2A, and three members of the TMEM16 family, xTMEM16A, xTMEM16E and xTMEM16K, are represented in the egg proteome (Figure 2). Second, we examined an RNA-seq time course in *X. laevis* oocytes and unfertilized eggs [27]. All four types of mRNA show increasing levels through gamete development, culminating in the egg (Figure 2). Although *ano6* and *clca3p-like* mRNA are present, it is likely that they are expressed after fertilization to guide the developing embryo through the maternal-to-zygotic transition, since their proteins are not detected in the unfertilized egg [34, 35].

**Figure 2.**
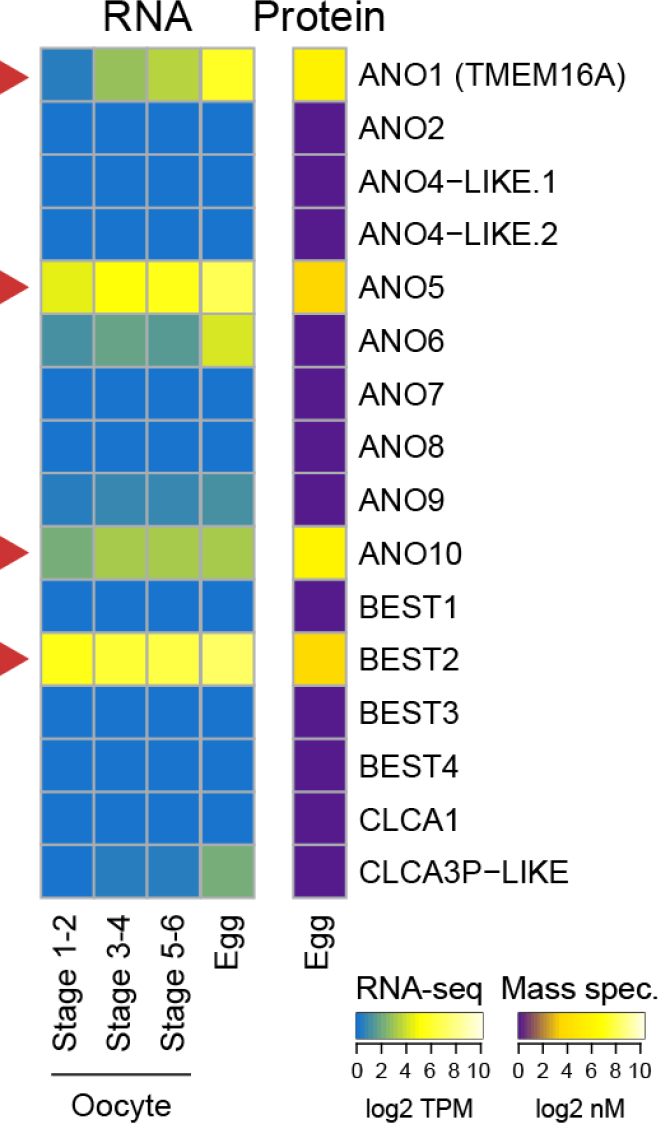
Expression of CaCCs in *X. laevis* oocytes and eggs. Heatmaps of expression levels CaCCs at the developmental stages indicated. (*Right*) Protein concentrations (from [28]) as determined by mass spectrometry-based proteomics study, in log_2_ nanomolar. (*Left*) Transcript levels (shown as transcripts per million (TPM), from [27]), as determined by RNA-seq-based transcriptome study). Red arrows highlight CaCCs with proteins found in eggs.

Both xBEST2A and xTMEM16A were originally cloned from fertilization-incompetent, *X. laevis* oocytes [21, 36], and each has been characterized as plasma membrane-localized [21, 36-38]. Moreover, xTMEM16A is the prominent CaCC in *X. laevis* oocytes [21]. In contrast, TMEM16E localizes to the endoplasmic reticulum (ER) where it functions as a Ca^2+^-activated scramblase [39, 40]. TMEM16K similarly localizes to the ER [41, 42]. Because both TMEM16E and TMEM16K proteins localize to the ER, they are not capable of passing the depolarizing Cl^−^ current of the fast block, and were therefore excluded both from further consideration. Together, these analyses suggest that the fast block to polyspermy in *X. laevis* eggs is mediated by either xBEST2A or xTMEM16A.

### Uncaging IP_3_ activates xTMEM16A and xBEST2A

Having discovered two CaCCs as candidates for the channel that mediates the fast block in *X. laevis* eggs, we sought to distinguish between their currents in the context of fertilization. Studying the activities of xTMEM16A and xBEST2A independently necessitated their exogenous expression. For this purpose, we chose a highly tractable system that lacks endogenous Ca^2+^-activated currents: *Ambystoma mexicanum (*axolotl) oocytes.

Although xTMEM16A was previously expressed in axolotl oocytes and currents generated in this context have been recorded [18], this is not the case for xBEST2A. We first confirmed that the exogenously expressed xBEST2A is localized to the plasma membrane of these oocytes. Confocal imaging of axolotl oocytes expressing both Ruby-tagged xBEST2A and the eGFP-tagged membrane marker MemE [43] revealed that xBEST2A was indeed expressed in these cells, and that it was transported to the plasma membrane (Figure 3A). As expected, no fluorescence was detected in water-injected control oocytes (Figure 3A).

**Figure 3.**
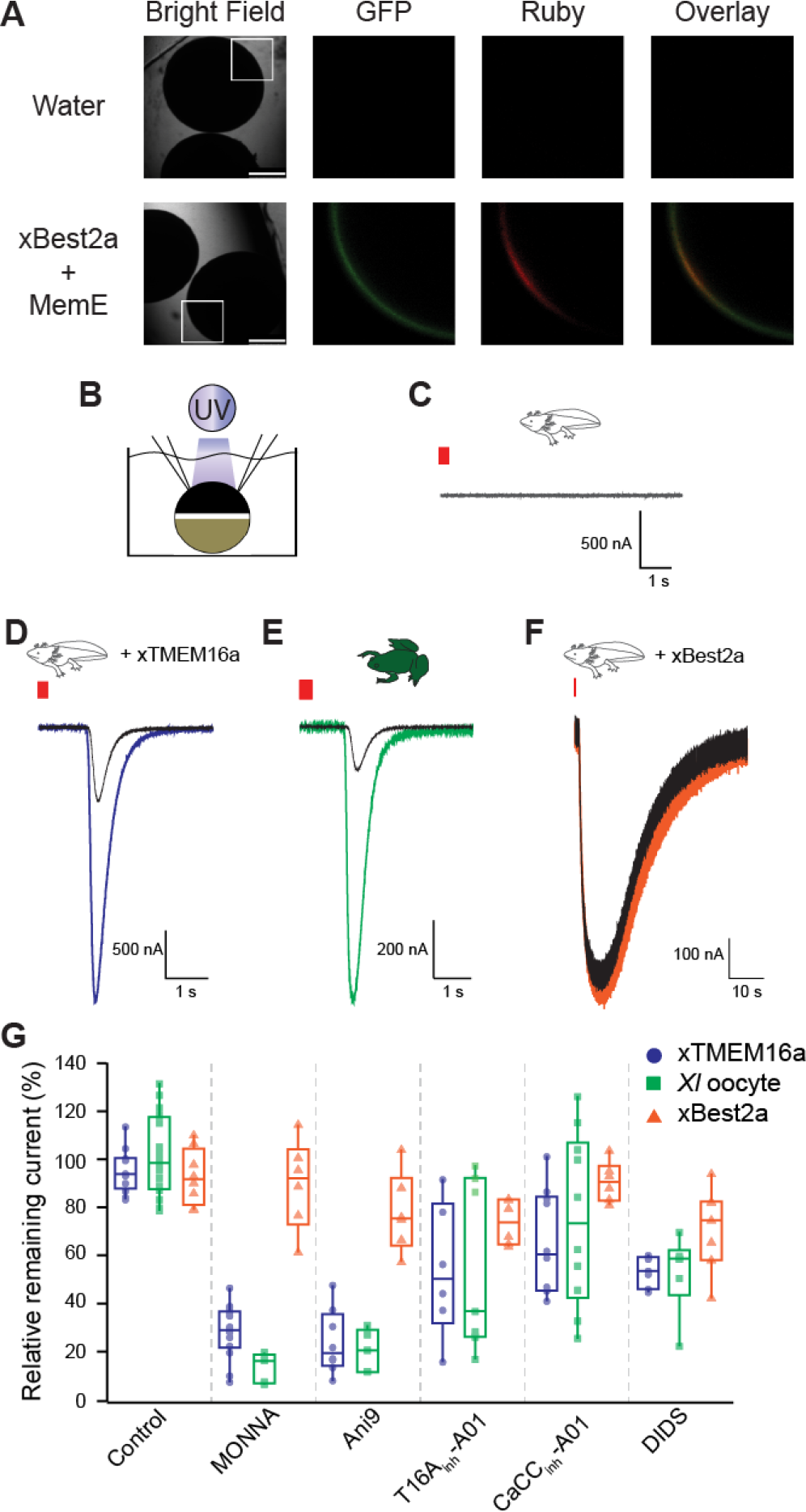
MONNA and Ani9 inhibit TMEM16A-conducted Cl^−^ currents. (A) Representative bright-field and fluorescence images of axolotl oocytes expressing Ruby-tagged xBEST2a and eGFP-tagged MemE (reporter of plasma membrane). Boxes denote portions included in fluorescence images, and scale bar denotes 750 (m. Overlay is of GFP and Ruby images. (B) Schematic of experimental design: UV photolysis to uncage IP_3_ while conducting TEVC. C-F) Current recordings from oocytes of (C-D *&* F) axolotls or (E) *X. laevis*, following injection with a photolabile caged IP_3_ analog, with clamping at −80 mV. Axolotl oocytes expressed (C) no transgene, (D) xTMEM16A, or (F) xBEST2A. (E) Wild-type *X. laevis* oocytes expressing endogenous channels. Typical current traces before and after uncaging, during (*colored*) control treatment and (*black*) in the presence of 10 μM MONNA. Red bar denotes the 250 ms duration of UV exposure. (*G*) Averaged proportion of current remaining after application of the indicated inhibitors, in axolotl oocytes expressing xTMEM16A (N=6-14) or xBEST2A (N=6-8), and in *X. laevis* oocytes expressing endogenous channels (N=5-16).

To study the currents conducted by xTMEM16A and xBEST2A, we exploited their shared regulation by Ca^2+^. Specifically, we photoactivated caged IP_3_ [21, 31, 32] by exposing the oocytes to ultraviolet light. This uncaging of IP_3_ induces Ca^2+^ release from the ER, thereby increasing the intracellular Ca^2+^ concentration and activating the channels (Figure 3B). As shown previously [21], uncaging IP_3_ in wildtype axolotl oocytes does not elicit any Ca^2+^-induced currents (Figure 3C). Importantly, we used the splice variants of xTMEM16A and xBEST2A channels that are present in *X. laevis* eggs [21, 36].

Using the uncaging system in conjunction with the two-electrode voltage clamp (TEVC), we recorded whole-cell currents in the presence or absence of known channel inhibitor molecules. Our initial assessment of the effects of sequential uncaging events in the absence of inhibitors revealed no differences in current between axolotl oocytes expressing either of the channels or *X. laevis* oocytes expressing the endogenous channels (Table S1). This finding indicated that differences in Ca^2+^-evoked currents measured in the presence or absence of an inhibitor in this system would reflect the efficacy of that inhibitor, thus this experimental design would enable us to characterize the efficacy of inhibitors in reducing xTMEM16A- or xBEST2A-mediated currents.

Using this set-up, we quantified the effects of five inhibitors on xTMEM16A- and xBest2a-mediated currents (Table S1). Three of these – MONNA, Ani9, and T16a_inh_-A01 – were previously reported to target human and/or mouse TMEM16A [44-46], whereas CaCC_inh_-A01 is a general inhibitor of CaCCs [44, 47]. Although no BEST-specific inhibitor has been characterized to date, we included the broad-spectrum Cl^−^ channel inhibitor DIDS because it reportedly binds to human bestrophin 1 (hBEST1) channels with an affinity 160-fold higher than that for mouse TMEM16A (mTMEM16A) [48].

### MONNA and Ani9 inhibit xTMEM16A currents

To characterize the effects of each inhibitor, we applied them to the above-described oocytes. In the case of xTMEM16A, both MONNA and Ani9 effectively reduced currents in the axolotl oocytes by over 70% (Figs. 3D & S2, Table S1), whereas T16A_inh_-A01 and CaCC_inh_-A01 were much less effective (Table S1, Figure S2). Unexpectedly, we found that 7.5 μM DIDS, a concentration well below the reported IC_50_ for the drug on mTMEM16A [48], reduced xTMEM16A by almost 50% (Table S1, Figure S2).

In *X. laevis* oocytes, the prominent Ca^2+^-activated Cl^−^ current is known to be generated by xTMEM16A channels [21]. Comparison of the effects on xTMEM16A-mediated current in the axolotl oocytes to the endogenous TMEM16A-passed currents generated in *X. laevis* oocytes revealed that in nearly all cases the efficacy of the inhibitors was very similar in the two test groups (Figure 3E & S2, Table S1). The exception is that MONNA blocked significantly more xTMEM16A current in the *X. laevis* oocyte (87 ± 2%) than in the axolotl oocytes (72 ± 3%) (*P*<0.05 ANOVA with post-hoc HSD Tukey; Table S1). Collectively, these data demonstrate that only MONNA and Ani9 effectively inhibit xTMEM16A.

### MONNA and Ani9 discriminate between currents generated by xTMEM16A and xBEST2A

Comparison of the effects of the five inhibitors on xBEST2A currents revealed that none had a significant effect (*P* > 0.05, ANOVA with post-hoc HSD Tukey; Figure 3F & S2 and Table S1). Most notably, currents generated in the presence of MONNA or Ani9 were no different than those produced in the control, confirming that these two compounds are specific for xTMEM16A. Furthermore, the lack of xBEST2A inhibition by MONNA and Ani9 demonstrates that these inhibitors do not interfere with the IP_3_-induced Ca^2+^ release pathway. Together, these results demonstrate that MONNA and Ani9 effectively target xTMEM16A channels but have only minimal effects on xBEST2A and the IP_3_ receptor, thereby providing a mechanism for discerning between xTMEM16A and xBEST2A currents during the fast block.

### The TMEM16A mediated-current produces the fast block in *X. laevis*

To characterize the fast block to polyspermy, we conducted whole-cell recording of *X. laevis* eggs during fertilization (Figure 4A). Eggs with steady resting potentials were inseminated with sperm and currents were recorded for up to 40 minutes or until the cortex contracted (indicating that fertilization was successful) (Figure 4B). Figure 4C depicts a typical fertilization-evoked depolarization that occurred after sperm addition. For eggs inseminated under control conditions, we found that: the resting potential was −19.2 ± 1.0 mV; the fertilization potential was 3.7 ± 2.3 mV (N=30, Figure 4D); the time between the addition of sperm and the onset of membrane depolarization (which likely represents the time required for the sperm to penetrate the viscous jelly coat of the egg [16]) was approximately 4.9 ± 0.7 minutes (N=30, Figure 4E); and the average rate of depolarization was 9.0 ± 3.4 mV/ms (N=30) (Figure 4F).

**Figure 4.**
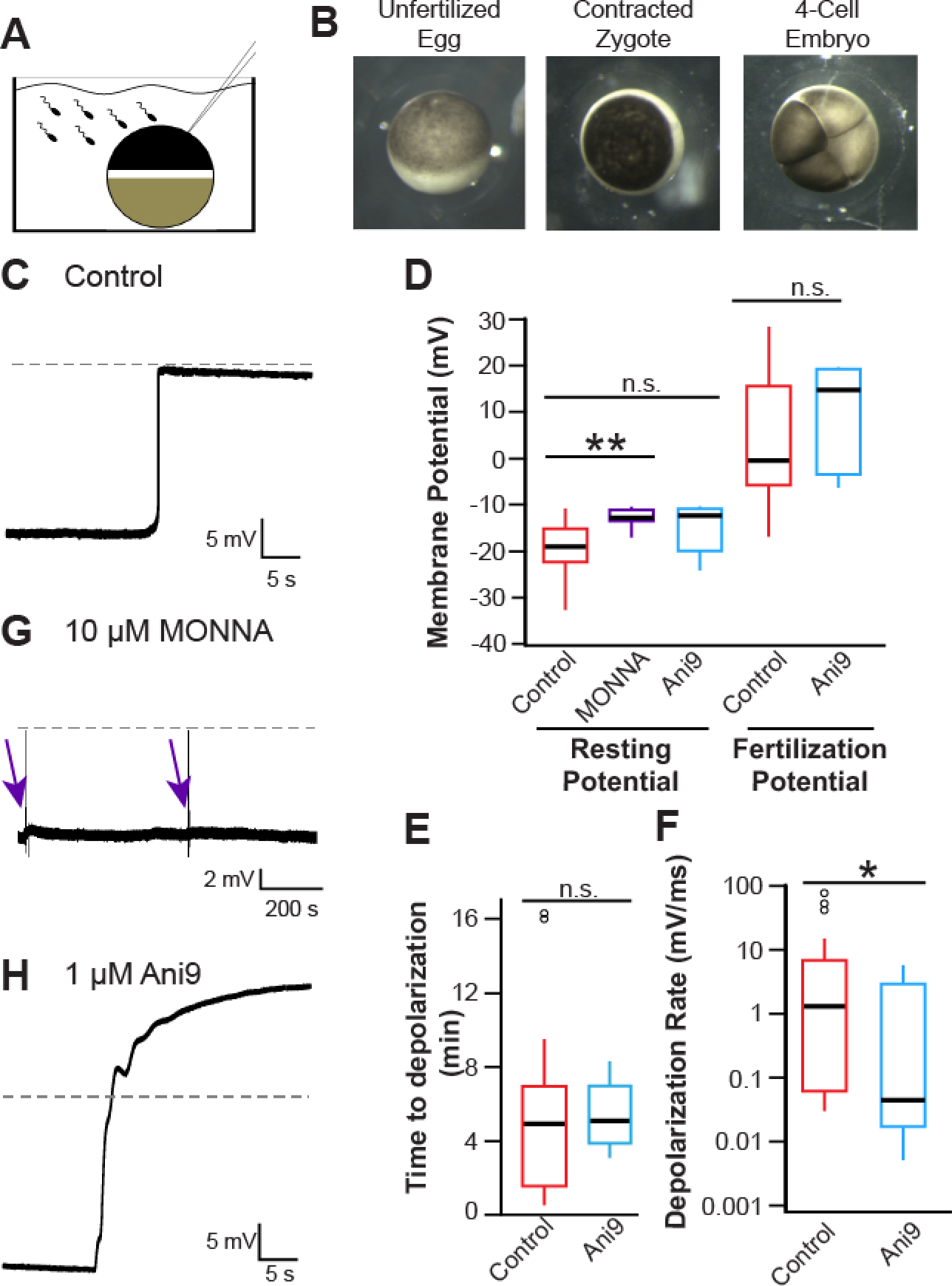
Fertilization activates TMEM16A to depolarize the egg. (*A*) Schematic depiction of experimental design: whole-cell recordings made on *X. laevis* eggs during fertilization. (*B*) Images of *X. laevis* (*left*) egg before sperm addition, (*center*) egg approximately 15 minutes after fertilization with animal pole contracted, and (*right*) 4-cell embryo. Representative whole-cell recordings made during fertilization in (*C*) control conditions, (G) the presence of 10 μM MONNA, (H) or the presence of 1 μM Ani9. Dashed lines denote 0 mV, purple arrows denote times at which sperm was applied to eggs in the presence of 10 μM MONNA. *D-F*) Tukey box plot distributions of (*D*) the resting and fertilization potentials in control conditions and with MONNA or Ani9, (*E*) the time between sperm application and depolarization in the absence and presence of Ani9, and (*F*) the depolarization rate in the absence and presence of Ani9 (N=5-30). ** denotes *P*<0.001, * denotes *P*<0.05, and n.s. denotes *P*>0.05.

To determine whether it is xTMEM16A or xBEST2A that conducts the depolarizing current responsible for the fast block, we inseminated eggs in the presence of MONNA or Ani9, each of which was expected to inhibit xTMEM16A but to have minimal effect on xBEST2a or IP_3_ receptors (Figure 3G, Table S1). In *X. laevis* eggs, inhibition of xTMEM16A using either inhibitor effectively diminished the fast block. In the presence of 10 μM MONNA, fertilization failed to evoke depolarization in seven independent experiments (Figure 4G); thus, this inhibitor completely abolished the fast block. Eggs incubated in MONNA had a significantly more positive resting potential than that of control eggs (−12.8 ± 0.8 mV vs. −19.2 ± 1.0 mV, T-test, *P* < 0.001) (Figure 4D). However, this elevated resting potential did not interfere with fertilization; visual assessment revealed contraction of the animal pole followed by the appearance of a cleavage furrows (Figure 4B), thus demonstrating that all eggs inseminated in the presence of MONNA initiated embryonic development.

In the presence of 1 μM Ani9, the rate of depolarization for inseminated eggs was significantly reduced, and thereby attenuating the fast block (1.2 ± 1.1 mV/ms with Ani9 (N=5) vs 9.0 ± 3.4 mV/ms in control (N=30), T-test, *P*<0.05) (Figure 4F & 4H). Because the rate of depolarization is proportional to the number of channels that are open, a slower rate reflects fewer channels being activated by fertilization. Based on the rates measured, we estimate that in the presence of 1 μM Ani9, 7.5-fold fewer channels were triggered to open by fertilization; i.e., only 13% of the channels that would be activated under normal conditions opened in this context. This 87% reduction in the number of open channels is consistent with the 80% inhibition of xTMEM1Aa channels measured when IP_3_ was uncaged in *X. laevis* and axolotl oocytes (Figure 3G, Table S1). No other metrics of the fast block differed significantly in recordings made in the presence vs. absence of Ani9 (Figure 4D – 4F).

Collectively, the inability of fertilization to depolarize an egg in the presence of MONNA and the slowed rate of depolarization in the presence of Ani9 demonstrate that TMEM16A channels produce the depolarizing current that mediates the fast block in *X. laevis* eggs.

## DISCUSSION

The fast block to polyspermy is one of the earliest and most prevalent events across species that undergo external fertilization. Despite its widespread use by evolutionarily divergent species, the signaling pathways that underlie these fertilization-evoked depolarizations have remained elusive. Here we identify the CaCC that mediates the fast block in the African clawed frog *X. laevis*: xTMEM16A (Figure 5). Given that an increase in the intracellular Ca^2+^ concentration and an efflux of Cl^−^ are required for the fast block in all frogs and toads studied thus far [7, 49], we propose that the current produced by TMEM16A channels triggers the fast block to polyspermy in all anurans.

**Figure 5.**
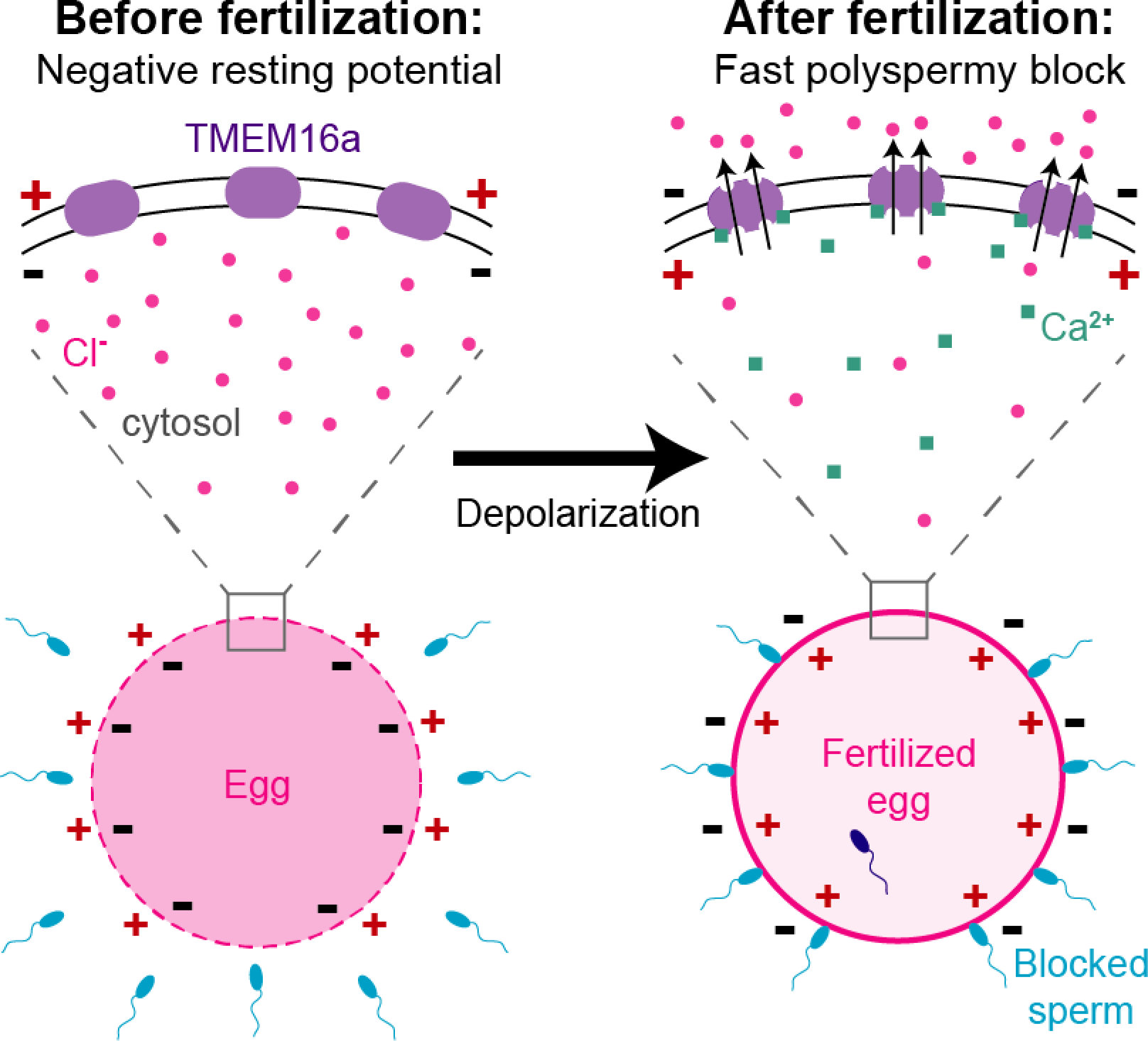
Proposed model for fertilization signaled activation of TMEM16A. Before fertilization, *X. laevis* eggs have a negative resting potential; thereby signaling to sperm that they can receive a male gamete. After fertilization, cytosolic Ca^2+^ increases to activate TMEM16a. An efflux of Cl^−^ then depolarizes the egg, and this change in membrane potential blocks supernumerary sperm from entering the fertilized egg.

Our identification of xTMEM16A and xBEST2A as candidate CaCCs that may mediate the fast block is based on proteomics and transcriptomics. Indeed, both proteins are translated in high concentrations (approximately 22 × 10^9^ xTMEM16A channels and 2 × 10^9^ xBEST2A channels, see Methods) in mature eggs. Given that these channels are present in the egg membrane, it was feasible that either or both could mediate the Ca^2+^-activated Cl^−^ efflux that drives the fast block in *X. laevis*.

Our finding that 10 μM MONNA and 1 μM Ani9, concentrations higher than their published IC_50_ [45, 46], inhibit >70% of xTMEM16A channels in both axolotl and *X. laevis* oocytes, yet that they are largely ineffective at reducing currents conducted by xBEST2A, strongly indicate that these inhibitors discriminate between our two candidate CaCCs. Both of these inhibitors are known to be highly specific for TMEM16A, with Ani9 failing to block even the closest relative of TMEM16A, TMEM16B [46]. In contrast, T16_inh_-A01, and CaCC_inh_-A01 were much less effective at inhibiting either xTMEM16A or xBEST2A. The similarity between the pharmacological profiles of xTMEM16A currents recorded in axolotl oocytes and endogenous Ca^2+^-activated currents in *X. laevis* oocytes supports the hypothesis that the native Ca^2+^-activated Cl^−^ currents in *X. laevis* oocytes are generated by xTMEM16A channels [21].

Although MONNA and Ani9 inhibited exogenously expressed xTMEM16A in axolotl oocytes to similar extents, MONNA was significantly more effective in reducing the endogenous Ca^2+^-activated currents of *X. laevis* oocytes (Figure 4F, *P* < 0.05, ANOVA with post-hoc HSD Tukey). The increased efficacy of MONNA with respect to endogenous xTMEM16A in the egg is consistent with the observed difference in its fertilization-induced electrical profile over that of Ani9 (i.e. with MONNA completely blocking depolarization and Ani9 merely slowing it). Given that the mechanisms underlying channel inhibition by these chemically distinct agents have not yet been elucidated, we hypothesize that the differing effects of these inhibitors on eggs, in spite of their similar effects on oocytes are attributable to the strikingly different environments at these two developmental time points. Furthermore, we speculate that the elevated resting potential recorded from eggs inseminated in the presence of MONNA reflects the altered Cl^−^ homeostasis in these cells, consistent with a recent demonstration that TMEM16A activity plays a prominent role in Cl^−^ homeostasis [46].

Previous studies showed that fertilization-evoked depolarization varies with respect to amplitude and shape, even when recorded under control conditions [7, 14, 17]. Our study further demonstrates that the rate of depolarization varies for each unique fertilization event. Because the depolarization rate is directly proportional to the number of channels that open, our data imply that different fertilization events lead to the opening of different numbers of channels. Although the source of the Ca^2+^ that signals the fast block remains to be determined, the variance in TMEM16A channel activation in response to fertilization may reflect variance in changes in Ca^2+^ levels between different eggs. For example, if fertilization triggers TMEM16A opening by a pathway that involves receptor activation and second-messenger signaling, variation may reveal that some sperm activate multiple receptors whereas others activate only one. By contrast, if fertilization stimulates Ca^2+^ entry to trigger the fast block, variation may be related to different numbers of Ca^2+^-permeant channels opening in response to fertilization. In other systems, TMEM16A can be activated by either receptor activated second messenger signaling or Ca^2+^ entry. For example, IP_3_-induced Ca^2+^ release activates TMEM16A in DRG neurons [38]; whereas, Ca^2+^ entry via TRPV6 channels activates TMEM16A in the epididymis [37].

Despite the gross changes that the plasma membrane of a *X. laevis* oocyte undergoes as it matures into a fertilization-competent egg, it is evident that the xTMEM16A channels are retained. Where in fertilization-competent eggs the xBEST2A channels localize remains to be determined. Based on its presence in the mature egg [28] and its lack of contribution to the fast block, we speculate that it is either desensitized or absent from the plasma membrane, as is the case for ORAI1 [23], PMCA [24], and Na^+^/K^+^ ATPase [25].

In conclusion, the fertilization-activated opening of TMEM16A channels is the earliest known signaling event evoked by the sperm-egg interaction (Figure 5). The discovery of a critical role for TMEM16A channels in fertilization lays a foundation for understanding how the membrane potential regulates fertilization. More broadly, TMEM16A channels regulate diverse processes ranging from epithelial secretions [30] to smooth muscle contraction [50, 51]. These CaCCs are indispensable for human health [52]. Due to their large size, ease and reproducibility for electrophysiology recordings, and years of study by developmental biologists and biophysicists alike, we propose that *X. laevis* fertilization may serve as a straightforward model system to study the physiologic regulation of this critically important channel.

## ACKNOWLEDGEMENTS

We thank B.L. Mayfield and E.R. Rochon for excellent technical assistance. We thank L.A. Jaffe and K.I. Kiselyov for helpful discussions and advice. This work was supported by an Andrew Mellon Predoctoral Fellowship to K.L.W., a March of Dimes Foundation Basil O’Connor Grant 5-FY16-307 to M.T.L., and NIH grant R00HD69410 to A.E.C.

## AUTHOR CONTRIBUTIONS

K.L.W., M.L., and A.E.C. conceived of the research. K.L.W., W.A.P., M.T., M.L., and A.E.C. created the experiments, designed their implementation, planned analyses, and wrote the manuscript.

## DECLARATION OF INTERESTS

The authors declare no competing interests.

## SUPPORTING INFORMATION

**FIGURE S1:**
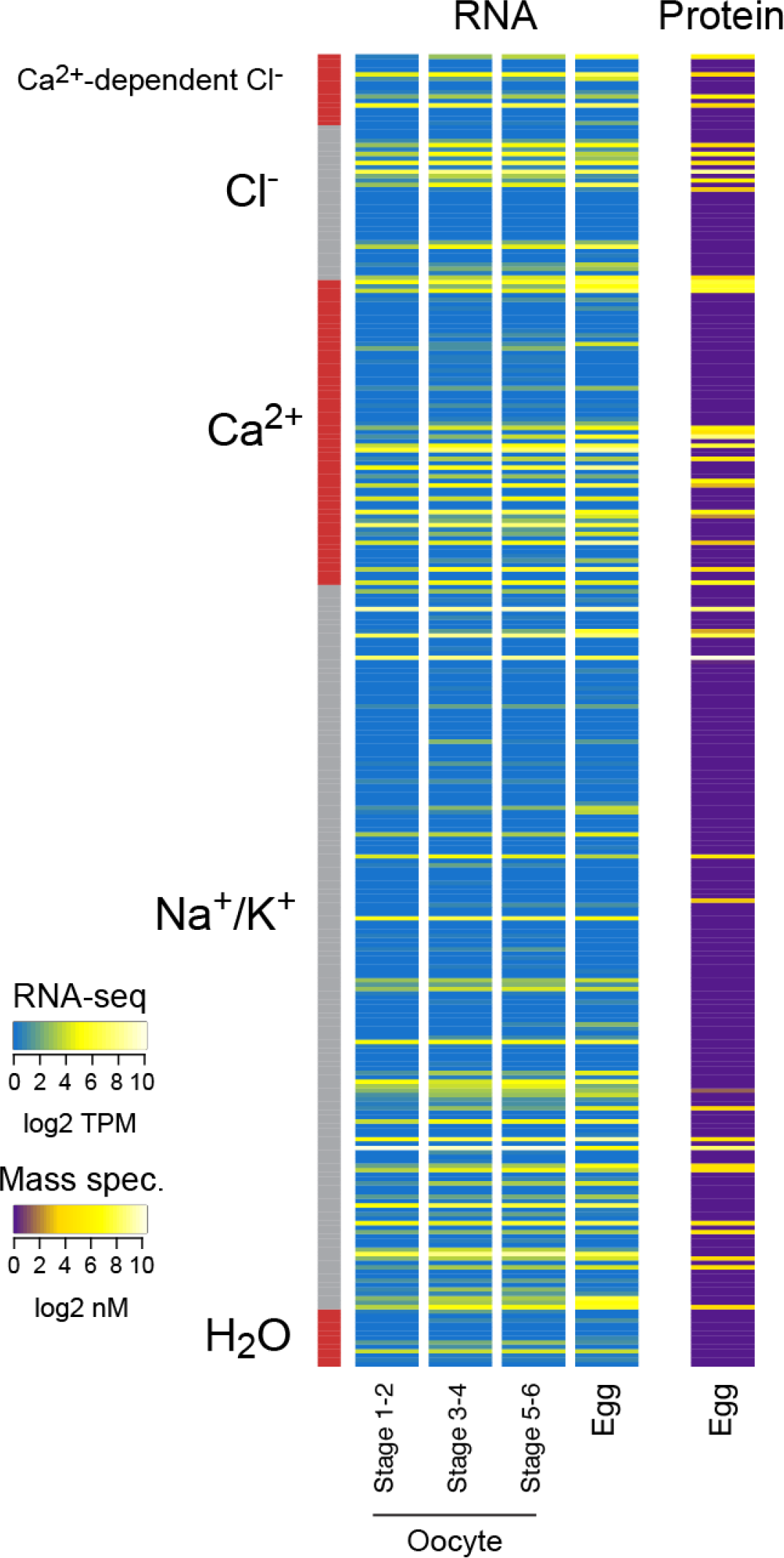
Heatmaps showing (*left*) RNA expression levels (based on RNA-seq from [27]), as log_2_ transcripts per million (TPM), and (*right*) protein concentrations (based on mass spectrometry from [28]) in log_2_ nanomolar. Transcripts and proteins are grouped by channel type.

**FIGURE S2:**
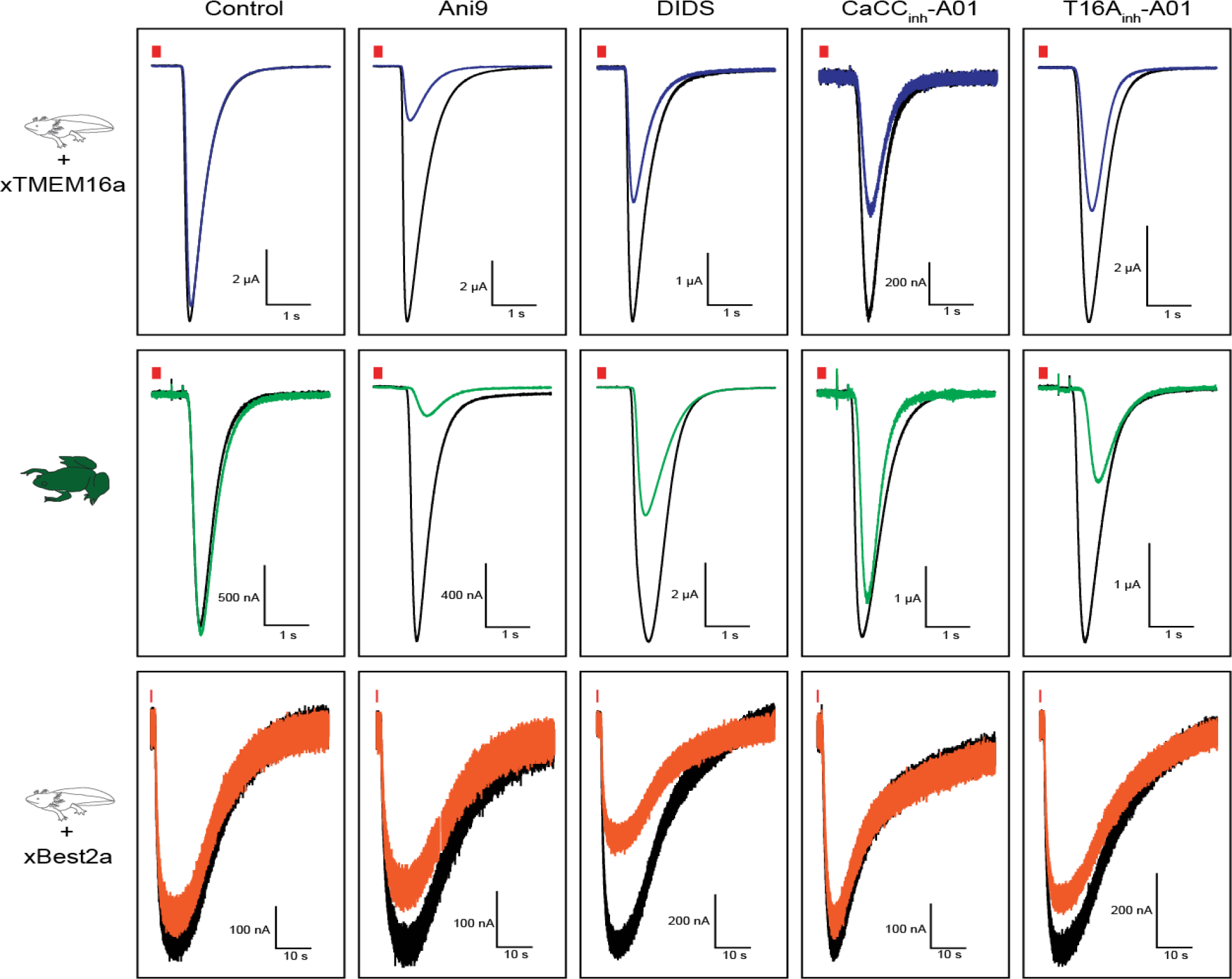
Representative current traces evoked by IP_3_ uncaging in axolotl oocytes expressing (*top*) xTMEM16A or (*bottom*) xBEST2A, and in (*middle*) wild-type *X*. *laevis* oocytes. Shown are typical traces (*black*) before and (*colored*) after application of a control solution, Ani9, DIDS, CaCC_inh_-A01, or T16A_inh_-A01. The red bars denote the 250 ms UV-exposure.

### Dataset S1

Gene Ontology terms used to identify channels; RNA-seq data from [27], of channels in *X. laevis* oocytes during developmental stages 1-2, 3-4, and 5-6, and in fertilization-competent eggs; and proteomics data from [28], from fertilization-competent *X. laevis* eggs.

**TABLE S1.** Inhibition of Ca^2+^-activated current using Cl^−^ channel inhibitors.

| | Control | 10 μM<br>MONNA | 1 μM<br>Ani9 | 30 μM<br>T16a<sub>inh</sub>-A01 | 10 μM<br>CaCC<sub>inh</sub>-A01 | 7.5 μM<br>DIDS |
| --- | --- | --- | --- | --- | --- | --- |
| **xTMEM16A**<br>**in axolotl**<br>**oocytes** | 5 ± 3 | 72 ± 3 | 76 ± 5 | 46 ± 11 | 35 ± 7 | 46 ± 2 |
| ***Xl* oocytes** | 2 ± 4 | 87 ± 2 | 80 ± 4 | 45 ± 13 | 25 ± 11 | 47 ± 7 |
| **xBEST2A**<br>**in axolotl**<br>**oocytes** | 7 ± 4 | 10 ± 8 | 22 ± 7 | 26 ± 4 | 9 ± 3 | 29 ± 7 |

Average ± SEM percentage (%) of current inhibition seen for uncaging experiments. The number of independent observations for each treatment is: MONNA (N=8-16); Ani9 (N=5-8); T16a<sub>inh</sub>-A01 (N=6-7); CaCC<sub>inh</sub>-A01 (N=6-10); DIDS (N=6-7). *Xl*: *Xenopus laevis*.

## METHODS

### Materials

N-((4-methoxy)-2-naphthyl)-5-nitroanthranilic acid (MONNA), 2-[(5-Ethyl-1,6-dihydro-4- methyl-6-oxo-2-pyrimidinyl)thio]-N-[4-(4-methoxyphenyl)-2-thiazolyl]-acetamide (T16A_inh_-A01), and 6-(1,1-Dimethylethyl)-2-[(2-furanylcarbonyl)amino]-4,5,6,7-tetrahydrobenzo[*b*]thiophene-3-carboxylic acid (CaCC_inh_-A01) were purchased from Sigma-Aldrich (St. Louis, MO), and 2-(4-chloro-2-methylphenoxy)-N-[(2-methoxyphenyl)methylideneamino]-acetamide (Ani9) from ChemDiv (San Diego, CA). Human chorionic gonadotropin (hCG) was purchased from Henry Schien (Melville, NY). All other materials, unless noted, were purchased from Thermo Fisher Scientific (Waltham, MA).

### Proteomic and RNA-seq analysis

Paired-end raw RNA-seq reads from [27] were downloaded from the NCBI Sequence Read Archive (SRA) (https://www.ncbi.nlm.nih.gov/sra) (accession numbers SRX1287719, SRX1287720, SRX1287721, and SRX1287707). Reads were aligned using HISAT2 [53] in paired-end mode with default parameters to the *X. laevis* v9.1 genome, obtained from Xenbase (http://www.xenbase.org), then assigned to genes using featureCounts [54] on Xenbase-annotated gene models in paired-end mode allowing multI-mappers (-p -M).

To identify channel genes, we assembled 106 relevant gene ontology (GO) terms that distinguished the following classes of channels: Cl^−^, Ca^2+^, Na^+^, K^+^, and H_2_O (Dataset S1). To account for possible gaps in GO term annotation, all family members of any gene annotated into a channel category were also included in further analysis; for example, all TMEM16 family members regardless of their GO annotation were included in this analysis.

To estimate the number of channels in the egg, we combined the protein concentrations with the stoichiometry of the functional channel: two subunits for TMEM16a channels [55] and five for Best2 [31, 56]. We then assumed that *X. laevis* eggs are spherical, and calculated their volume based on their measured diameter of 1.4 mm [57].

### Solutions

#### Fertilization solutions

Modified Ringers (MR) (in mM): 100 NaCl, 1.8 KCl, 2.0 CaCl_2_, 1.0 MgCl_2_, and 5.0 HEPES, pH 7.8, and filtered using a sterile, 0.2 μm polystyrene filter [58]. Fertilization recordings were made in our standard solution of 20% MR (also known as MR/5) with or without inhibitors, as indicated. After electrical recordings were made for fertilization experiments, embryos developed for two hours in 33% MR (MR/3). Various recordings were made in the presence of inhibitors, either diluted in water or ≤ 2% dimethyl sulfoxide (DMSO).

#### Oocyte solutions

Oocyte Ringers 2 (OR2) (in mM): 82.5 NaCl, 2.5 KCl, 1 MgCl_2_, and 5 mM HEPES, pH 7.6 [59].

#### Two-electrode voltage clamp solution

ND96 (in mM): 96 NaCl, 2 KCl, 1 MgCl_2_, 10 HEPES, pH 7.6 and filtered with a sterile, 0.2 μm polystyrene filter [21].

### Animals

*Xenopus laevis* adults were obtained commercially (RRID: NXR_0.0031, NASCO, Fort Atkinson, WI), as were axolotls, *Ambystoma mexicanum* (RRID: AGSC_100A, Ambystoma Genetic Stock Center, Lexington, KY). *X. laevis* and axolotls were housed separately at 18 °C with 12/12-hour light/dark cycle.

### Collection of Gametes, Fertilization, and Developmental Assays

All animal procedures were conducted using accepted standards of humane animal care and were approved by the Animal Care and Use Committee at the University of Pittsburgh.

*X. laevis* and axolotl oocytes were collected by procedures similar to those described previously [21, 57]. Briefly, ovarian sacs were obtained from *X. laevis* females anesthetized with a 30-minute immersion in 1.0 g/L tricaine-S (MS-222) at pH 7.4 and axolotls euthanized via immersion in 3.6 g/L tricaine-S at pH 7.4. For both sets of oocytes, ovarian sacs were manually pulled apart and incubated for 90 minutes in 1 mg/ml collagenase in ND96 supplemented with 5 mM sodium pyruvate and 10 mg/L of gentamycin. Collagenase was removed by repeated washes with OR2, and healthy oocytes were sorted and stored at 14 °C in ND96 with sodium pyruvate and gentamycin.

Eggs were collected from sexually mature *X. laevis* females as previously described [57]. Briefly, females were injected 1,000 IU of hCG into their dorsal lymph sac and housed overnight for 12-16 hours at 14-16 °C. Typically, females began laying eggs within 2 hours of moving to room temperature. Eggs were collected on dry petri dishes and used within 10 minutes of being laid.

Sperm were harvested from testes of sexually mature *X. laevis* males, as previously described [57]. Following euthanasia by a 30-minute immersion in 3.6 g/L tricaine-S (pH 7.4), testes were dissected. Cleaned testes were stored at 4 °C in MR for usage on the day of dissection or in L-15 medium for use up to one week later.

To create a sperm suspension, approximately 1/10 of a testis was minced in MR/5; if not used immediately, this solution was stored on ice and used for up to one hour. No more than three sperm additions were added to a given egg during whole cell recordings, and the total volume of sperm suspension added never exceeded 7.5% of the total fertilization solution. Eggs inseminated during whole cell recordings were transferred to MR/3 for up to 47 hours after insemination to monitor development. Development was assessed based on the appearance of cleavage furrows (Figure 4B), which were typically apparent approximately 90 minutes after sperm addition [57].

### Electrophysiology

Electrophysiology recordings were made using TEV-200A amplifiers (Dagan Co.) and digitized by Axon Digidata 1550A (Molecular Devices). Data were acquired with pClamp Software (Molecular Devices) at a rate of 5 kHz.

IP_3_-evoked currents were recorded in the two-electrode voltage clamp (TEVC) configuration at −80 mV, from *X. laevis* or axolotl oocytes. The cDNA encoding the *X. laevis* xTMEM16A channel in the GEMHE vector was provided by L. Jan (University of California San Francisco) [21]. The cDNA encoding the xBEST2A channel was purchased from DNASU [60] and was engineered into the GEMHE vector with a carboxy-terminal Ruby tag [61] using overlapping PCR and Gibson assembly methods. The sequences for all constructs were verified by automated Sanger sequencing (Gene Wiz). The xTMEM16A and xBEST2A cRNAs were transcribed using the T7 mMessage mMachine Ultra kit (Ambion), and MemE with the SP6 mMessage mMachine kit (Ambion). Defolliculated axolotl oocytes were injected with 5 ng of cRNA for xTMEM16a or xBest2a, as described previously [21]. Both axolotl and *X. laevis* oocytes were injected with the photolabile IP_3_ analog: *myo*-inositol 1,4,5-trisphosphate, *P*4(5)-1-(2-nitrophenyl) ethyl ester (caged-IP_3_). Each oocyte was injected with a 200 μM caged-IP_3_ stock made in DDH_2_O to reach a final concentration of 5 μM within the oocyte [21], and incubated in the dark at 18 °C for 1-5 hours before recording. Pipettes of 1-8 MΩ resistance were pulled from borosilicate glass and filled with 1 M KCl. The nitrophenyl cage on IP_3_ was released by flash photolysis with a 250 ms exposure to light derived from the Ultra High Power White LED Illuminator (380-603 nm, Prizmatix) and guided by a liquid light source to the top of oocytes in our recording chambers (RC-26G, Warner Instruments). Using the TEVC technique, we recorded Ca^2+^-activated Cl^−^ currents ranging from 0.2 to 17 μA, with an average of 6.9 ± 1.5 μA in *X. laevis* oocytes (N=16), 5.6 ± 1.0 μA (N=12) for xTMEM16a in axolotl oocytes, and 0.47 ± 0.7 μA for xBest2a in axolotl oocytes. The bath solution was changed with the gravity fed, pinch valve VC-8 solution changer (Warner Instruments). Background-subtracted peak currents were quantified from two consecutive recordings: one before and one with application of the tested inhibitors. The proportional difference between peak currents before and with inhibitor application for each oocyte was used to quantify the percent inhibition for each treatment. It is not possible to compare current amplitudes generated in different oocytes directly due to the innate variability of the experimental set-up (*i.e*. positioning of the UV light, exact amount of caged IP_3_ in each oocyte, etc.).

Fertilization-evoked depolarizations were recorded in the whole cell configuration. Pipettes of 5-20 MΩ resistance were pulled from borosilicate glass and filled with 1 M KCl. Resting and fertilization potentials were generally stable and quantified approximately 10 seconds before and after the depolarization, respectively. Depolarization rates were quantified by determining the maximum velocity of the quickest 1 mV shift in the membrane potential for each recording.

### Imaging

Axolotl oocytes were imaged using a TCS SP5 confocal microscope (Leica Microsystems, Wetzlar, Germany) equipped with a Leica 506224 5X objective. As a membrane control, oocytes were injected with cRNA for a membrane anchored eGFP [43] which was *in vitro* transcribed using an SP6 mMessage mMachine kit (Ambion). EGFP was excited with a 488 nm visible laser, whereas Ruby was excited with a 561 nm laser. Using a galvo scanner with unidirectional (600 Hz) scanning, sequential frames were captured with 2x line averaging. Images were analyzed using LAS AF (version 3.0.0 build 834) software and ImageJ [62].

### Quantification and Statistical Analyses

All electrophysiology recordings were analyzed with Igor (WaveMetrics) and Excel (Microsoft). Averaged values ± standard error of the means (SEM), are reported for each experimental condition. T-tests (one-tailed for depolarization rates and two-tailed for resting and fertilization potentials and comparisons of relative amplitudes of IP_3_-evoked currents) were used to determine differences between inhibitor treatments. Depolarization rates were log10 transformed before statistical analysis. ANOVAs followed by post-hoc HSF Tukey tests were used to compare different currents recorded with the same inhibitors.

